# 16p12.1 deletion orthologs are expressed in motile neural crest cells and are important for regulating craniofacial development in *Xenopus laevis*

**DOI:** 10.1101/2020.12.11.421347

**Authors:** Micaela Lasser, Jessica Bolduc, Luke Murphy, Caroline O’Brien, Sangmook Lee, Santhosh Girirajan, Laura Anne Lowery

## Abstract

Copy number variants (CNVs) associated with neurodevelopmental disorders are characterized by extensive phenotypic heterogeneity. In particular, one CNV was identified in a subset of children clinically diagnosed with intellectual disabilities (ID) that results in a hemizygous deletion of multiple genes at chromosome 16p12.1. In addition to ID, individuals with this deletion display a variety of symptoms including microcephaly, seizures, cardiac defects, and growth retardation. Moreover, patients also manifest severe craniofacial abnormalities, such as micrognathia, cartilage malformation of the ears and nose, and facial asymmetries; however, the function of the genes within the 16p12.1 region have not been studied in the context of vertebrate craniofacial development. The craniofacial tissues affected in patients with this deletion all derive from the same embryonic precursor, the cranial neural crest, leading to the hypothesis that one or more of the 16p12.1 genes may be involved in regulating neural crest cell (NCC)-related processes. To examine this, we characterized the developmental role of the 16p12.1-affected gene orthologs, *polr3e*, *mosmo*, *uqcrc2*, and *cdr2*, during craniofacial morphogenesis in the vertebrate model system, *Xenopus laevis*. While the currently-known cellular functions of these genes are diverse, we find that they share similar expression patterns along the neural tube, pharyngeal arches, and later craniofacial structures. As these genes show co-expression in the pharyngeal arches where NCCs reside, we sought to elucidate the effect of individual gene depletion on craniofacial development and NCC migration. We find that reduction of several 16p12.1 genes significantly disrupts craniofacial and cartilage formation, pharyngeal arch migration, as well as NCC specification and motility. Thus, we have determined that some of these genes play an essential role during vertebrate craniofacial patterning by regulating specific processes during NCC development, which may be an underlying mechanism contributing to the craniofacial defects associated with the 16p12.1 deletion.

## Introduction

Embryonic development requires the proper function of thousands of genes in order for cells to proliferate and divide, differentiate, migrate long distances to their final destination, and communicate with one another appropriately. Disruption of protein function due to mutations of genes that are required for these processes during embryogenesis can lead to severe developmental defects and neurodevelopmental disorders such as intellectual disabilities (ID) or Autism spectrum disorder (ASD) (Alonso-Gonzalez et al., 2018; Chen et al., 2019; Kasherman et al., 2020; Lasser et al., 2018; Sierra-Arregui et al., 2020). Genetic mutations caused by rare copy number variants (CNVs), including deletions and duplications, have been associated with neurodevelopmental disorders to varying degrees (Blazejewski et al., 2018; Deshpande and Weiss, 2018; Jensen et al., 2018; Pizzo et al., 2019; Rylaarsdam and Guemez-Gamboa, 2019; Singh et al., 2020). Recently, a CNV was identified in children diagnosed with ID, that results in a hemizygous deletion of several genes located at chromosome 16p12.1 (Antonacci et al., 2010; Girirajan et al., 2010). Individuals with this variant display phenotypic variability, presenting with a wide range of defects including microcephaly, seizures, cardiac defects, and growth retardation. Additionally, patients manifest severe craniofacial abnormalities including facial asymmetries, micrognathia (undersized jaw), a short philtrum (space between the nose and lip), as well as cartilage malformation of the ears and nose.

Craniofacial defects are one of the most prevalent congenital defects that can severely affect quality of life (Kirby, 2017; Trainor, 2010; Vega-Lopez et al., 2018). As craniofacial patterning relies heavily on the specification, proliferation, and subsequent migration of neural crest cells (NCCs), many craniofacial and cartilage defects arise due to aberrant NCC development (Fish, 2016; Rutherford and Lowery, 2016; Trainor, 2010; Vega-Lopez et al., 2018). Several of the tissue and organ systems affected by the 16p12.1 deletion are derived from NCCs; however, the function of the genes within this region have not been investigated in the context of vertebrate craniofacial development, nor has any study determined whether their depletion might impact NCC behavior. Therefore, the underlying developmental mechanism by which each gene contributes to the craniofacial phenotypes associated with the deletion remains to be elucidated.

The multigenic nature of the 16p12.1 deletion adds complexity to our understanding of the etiology behind this variant, due in part to the functional diversity of the affected genes. The 16p12.1 deletion encompasses a 520-kb region on the short arm of chromosome 16 and impacts several genes including *POLR3E*, *MOSMO*, *UQCRC2*, and *CDR2* (Antonacci et al., 2010; Girirajan et al., 2010). RNA polymerase III subunit E (POLR3E) is important for catalyzing the transcription of small RNAs, though the exact function of *POLR3E* in relation to RNA polymerase III activity is unknown (Hu et al., 2002). Modulator of smoothened (*MOSMO*) is a negative regulator of Sonic Hedgehog (Shh) signaling by participating in the degradation of the Frizzled Class receptor, Smoothened (Pusapati et al., 2018). Additionally, reduced dosage of *polr3e* and *mosmo* were found to severely impact proper brain development in both *Drosophila melanogaster* and *Xenopus laevis*, suggesting that they may contribute to the ID and microcephaly phenotypes observed in patients with the 16p12.1 deletion (Pizzo et al., 2020). Ubiquinol-cytochrome C reductase core protein 2 (*UQCRC2*) is a component of the mitochondrial respiratory chain complex and is essential for the production of ATP (Gaignard et al., 2017; Miyake et al., 2013; Shan et al., 2019; Shang et al., 2018). Cerebellar degeneration related protein 2 (*CDR2*) is an onconeural protein that is known to be ectopically expressed in breast or ovarian tumors, resulting in the generation of autoantibodies that leads to cerebellar degeneration (Hwang et al., 2016; Schubert et al., 2014).

Due to the broad range of cellular functions of each 16p12.1-affected gene, it is imperative to determine whether their individual depletion leads to specific craniofacial defects, or whether depletion of multiple genes within this region combinatorially contribute to a collaborative craniofacial phenotype. Thus, we investigated the contributions of *polr3e*, *mosmo*, *uqcrc2*, and *cdr2* to developmental processes that govern early craniofacial patterning in *Xenopus laevis*. First, we examined expression profiles for each transcript across early stages of embryonic development, and we observed their expression in motile NCCs residing in the pharyngeal arches (PAs), suggesting that they may influence NCC development and migration. Knockdown (KD) strategies were then utilized to assess the contribution of each 16p12.1-affected gene to facial and cartilage development. We previously showed that depletion of *polr3e*, *mosmo*, and *uqcrc2* led to smaller facial features (Pizzo et al., 2020), and in the present study, we find that these three genes also severely disrupt cartilage morphology. We then performed both *in vivo* and *in vitro* NCC migration assays, observing that some of these gene mutations also directly impact NCC migration in the pharyngeal arches and NCC motility rates. Finally, we examined NCC specification and proliferation, and while we found that reduced dosage of each gene did not have a significant impact on proliferation, we found that some of these genes are critical for NCC specification. Together, our results support the hypothesis that the craniofacial phenotypes associated with the 16p12.1 deletion are, in part, due to several genes within this region performing critical functions during NCC development and migration, craniofacial patterning, and cartilaginous tissue formation. Moreover, this work is the first to elucidate the roles of the 16p12.1-affected genes during embryonic craniofacial morphogenesis on a shared, directly-comparable background, providing deeper insight into how diverse genetic mutations lead to distinct developmental phenotypes and disease within the context of a multigenic syndrome.

## Results

### 16p12.1-affected genes display expression in the developing nervous system, pharyngeal arches, and craniofacial structures

One of the more prominent symptoms in patients with the 16p12.1 deletion are craniofacial dysmorphisms with varying severity (Girirajan et al., 2010). Children often present with facial asymmetries, micrognathia, a short philtrum, small and deep-set eyes, hypertelorism, a depressed nasal bridge, and dysplastic ears. Comorbidities commonly include microcephaly, growth retardation, scoliosis, and defects in hand and foot development (Girirajan et al., 2010). As proper NCC specification, proliferation, and migration are critical for governing embryonic facial patterning, we hypothesized that one or more of the 16p12.1-affected genes are required for NCC development, and that their depletion would result in defects associated with one or more of these NCC-related processes.

First, we investigated the spatiotemporal expression of four 16p12.1-affected gene orthologs, *polr3e*, *mosmo*, *uqcrc2*, and *cdr2*, across multiple stages of development in *Xenopus laevis* embryos. To examine this, we performed whole-mount *in situ* hybridization with DIG-labeled antisense RNA probes against these four genes (**Figure 1** and **Figure S1**; for *in situ* hybridization controls against RNA sense strands, see **Figure S2**). Interestingly, we observed primarily ubiquitous expression of all four gene transcripts during early blastula and gastrula stages (st. 10 and 13; data not shown; st. 20; **Figures S1A-B,E-F,I-J,M-N**). However, by stage 25, more defined expression became visible during early craniofacial morphogenesis with expression of each gene in migratory NCCs that reside in the pharyngeal arches (**Figures 1B-E**), and their expression patterns closely resemble the NCC-enriched transcription factor, *twist* (**Figure 1A**). At this stage, expression of both *polr3e* and *mosmo* is particularly strong in the developing brain and eye, which is consistent with previous data suggesting that these genes are important for both brain and eye development (Pizzo et al., 2020). All four genes are expressed in the developing head and facial structures throughout stage 35 (**Figures S1C-D,G-H,K-L,O-P**); *polr3e* expression appears to become more defined in the hindbrain region, whereas *mosmo* expression is stronger in the forebrain region (**Figures S1C,G**), and *uqcrc2* expression is heavily expressed in the developing kidney and somites (**Figure S1K**). By stage 40, *mosmo* expression is also observed in the developing spinal cord (**Figure S1H**). Additionally, the expression patterns of all four genes show potential overlap with cardiac tissue (**Figures S1D,H,L,P**). Thus, our findings demonstrate that the four 16p12.1-gene orthologs display expression in the developing nervous system, migratory NCCs in the pharyngeal arches, and later craniofacial structures, among other tissues.

**Figure 1:**
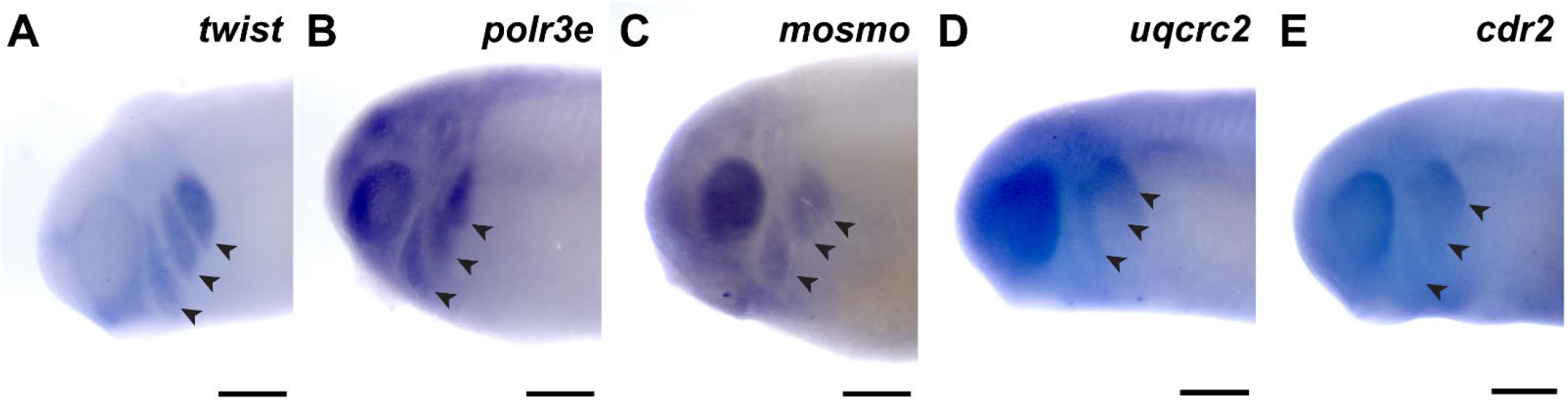
16p12.1-affected genes are expressed in migrating neural crest cells during embryonic development. **(A)** Lateral view of whole-mount *in situ* hybridization at stage 25 for *twist*, a NCC-enriched transcription factor. Anterior of embryo is to left. Arrows indicate the pharyngeal arches (PAs). **(B-E)** *In situ* hybridization at stage 25-28 for *polr3e*, *mosmo*, *uqcrc2*, and *cdr2* demonstrate expression in NCCs that occupy the PAs (n = 10 per probe). Scalebar = 300μm.

### Several 16p12.1-affected genes are important for maintaining cartilage size and scaling

Many individuals with the 16p12.1 deletion display defects in cartilage and skeletal development including deformed nose and ears, tooth malformation, short stature, smaller head size, and delayed growth (Girirajan et al., 2010). Additionally, patients also have speech, feeding, and swallowing impairments that are linked to abnormal jaw and throat formation (Girirajan et al., 2010). As these cartilaginous and skeletal tissues are derived from NCCs (Etchevers et al., 2019; Merkuri and Fish, 2019; Szabo and Mayor, 2018; Van Otterloo et al., 2016), we hypothesized that one or more of the 16p12.1-affected genes may play an essential role during embryonic development of craniofacial cartilage and skeletal structures. To examine this, we performed partial depletion of the four 16p12.1 gene orthologs to determine their influence on cartilage scaling and morphology in *Xenopus laevis* embryos (**Figure 2**). As the human disorder is due to a hemizygous deletion, we sought to achieve 50% knockdown in an attempt to recapitulate the gene dosage of the human condition. Thus, we utilized morpholino antisense oligonucleotides, which we previously validated to achieve 50% knockdown for each gene (Pizzo et al., 2020).

**Figure 2:**
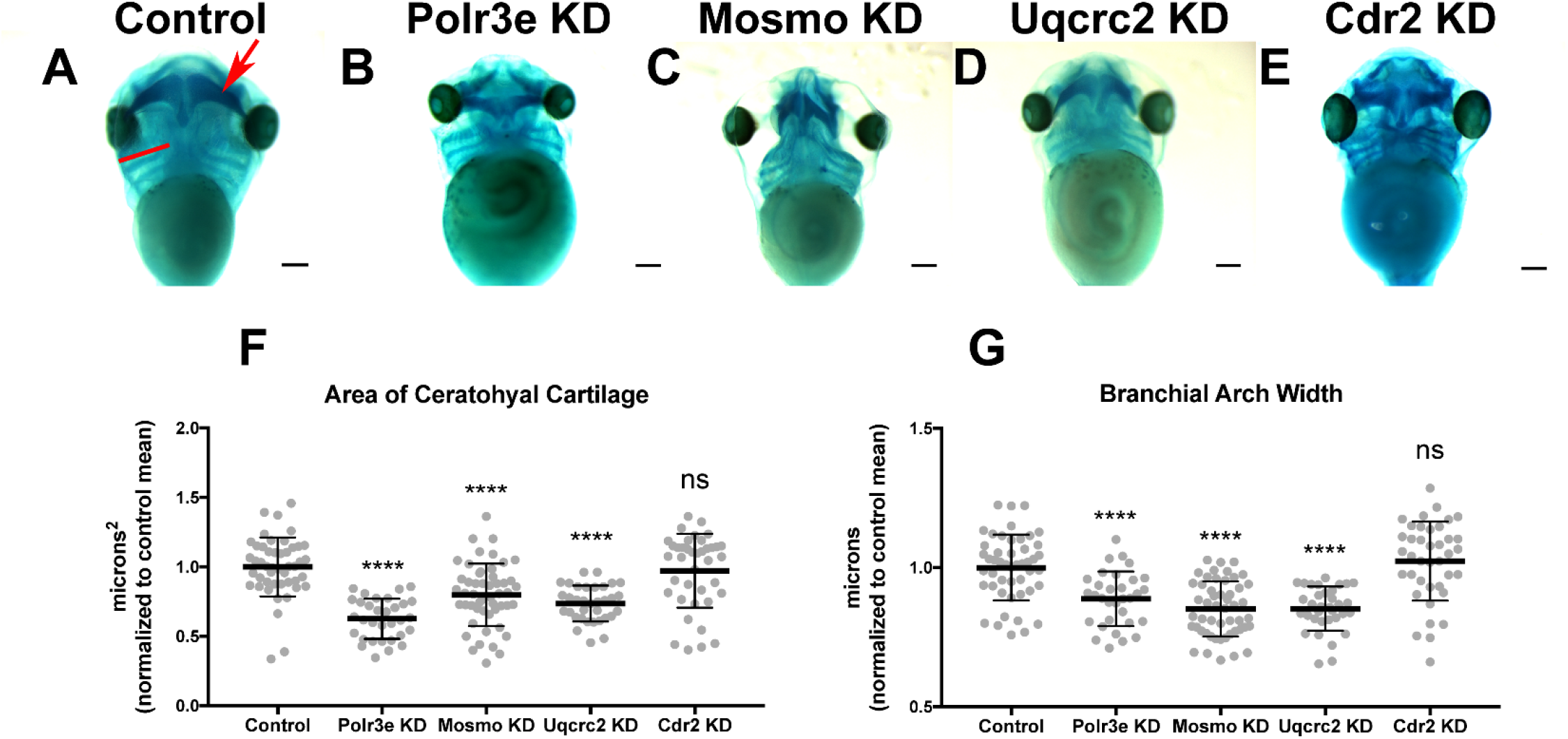
Knockdown of Polr3e, Mosmo, and Uqcrc2 impact cartilage morphology. **(A-E)** Ventral view of stage 42 embryos following single 16p12.1-associated gene KD, stained with Alcian blue to label cartilage elements. Anterior of embryo is at the top. In the control, the red arrow points to ceratohyal cartilage, while the red bar spans the first branchial arch. **(F-G)** Measurements of the average ceratohyal area and width of the first branchial arch. The data was normalized to the control MO condition. Partial depletion of Polr3e, Mosmo, and Uqcrc2 significantly reduced the size of both of these cartilage elements compared to controls, while depletion of Cdr2 had no effect on cartilage size. Significance determined using a student’s unpaired *t*-test. (Embryos quantified: Control = 48, Polr3e KD = 32, Mosmo KD = 51, Uqcrc2 KD = 34, Cdr2 KD = 39). \*\*\*\**p* < 0.0001, \*\*\**p* < 0.001, \*\**p* < 0.01, \**p* < 0.05, n.s., not significant. Scalebar = 300μm.

Reduced dosage of *polr3e*, *mosmo*, and *uqcrc2* was sufficient to severely impact cartilage development of stage 42 embryos. When we knocked down these three genes individually (throughout the entire embryo), we observed overt defects on cartilage morphology compared to controls (**Figures 2B-D**), including a decrease in the average ceratohyal area and the width of the first branchial arch (**Figures 2F-G**). Surprisingly, given that we previously showed that partial depletion of *cdr2* led to a minor reduction in facial size (Pizzo et al., 2020), this did not cause similar overt cartilage abnormalities (**Figure 2E**), nor was there any obvious effect on the size of individual cartilage elements (**Figures 2F-G**). However, the craniofacial defects associated with reduced dosage of *cdr2* are quite mild, and thus, this gene may not play a major role during craniofacial and cartilage morphogenesis when partially depleted, compared to the other genes affected in the 16p12.1 region. It is also possible that there may be genetic compensation by other closely related genes such as *cdr1*, *cdr2-like*, and *cdr3*. Overall, these results indicate that Polr3e, Mosmo, and Uqcrc2 are essential for early cartilaginous tissue formation and are likely important for later development of head and facial skeletal structures. Moreover, these results demonstrate that partial depletion of the 16p12.1-associated genes creates persistent defects on craniofacial patterning and cartilage formation that are not ameliorated later in development (stage 42), leading to our hypothesis that these genes may impact processes important for the embryonic progenitors of these tissues.

### Several 16p12.1-affected genes are critical for normal pharyngeal arch migration and NCC motility

Given that the 16p12.1-affected gene transcripts display expression in NCCs residing in the pharyngeal arches during stages that correspond with their migration (st. 25-30), we hypothesized that depletion of one or more of these genes may disrupt NCC migration and motility. To test this, we utilized single-blastomere injection strategies to generate left-right chimeric embryos, allowing for a side-by-side comparison of *twist* expression patterns between wild-type or KD sides, and we tracked the progress of migratory NCCs (**Figure 3**). Following single-sided individual depletion of each 16p12.1 gene, embryos were staged to 25-28, fixed, and *in situ* hybridization was performed against *twist*. To quantify NCC migration away from the anterior neural tube, measurements were taken of the total pharyngeal arch (PA) area, length of each individual pharyngeal arch, and total migration distance of the NCCs that form each individual pharyngeal arch, for both the uninjected and control or KD side of each embryo (**Figures 3F-Q**). We found that Polr3e and Mosmo knockdown significantly reduced total area of NCC streams (**Figures 3F-G**). Further, when Polr3e levels were reduced, the posterior PA was shorter in length (**Figure 3J**), whereas knockdown of Mosmo reduced the length of the anterior PA and hyoid PA (**Figure 3L**). Additionally, individual depletion of these genes reduced the ventral migration distance of all three NCC streams compared to controls (**Figures 3K,M**). Interestingly, knockdown of Uqcrc2 resulted in an increase in total PA area (**Figure 3H**) and a slight increase in the length of the anterior and hyoid PAs (**Figure 3N**), though it did not impact NCC migration (**Figure 3O**). It is possible that NCC proliferation is upregulated to compensate for reduced Uqcrc2 levels, leading to an increase in PA area and length. While knockdown of Cdr2 slightly increased hyoid PA length (**Figure 3P**), it did not result in significant changes in PA area or NCC migration (**Figure 3I,Q**). Together, these results suggest a specific role for Polr3e, Mosmo, and Uqcrc2 in maintaining NCC migration *in vivo*.

**Figure 3:**
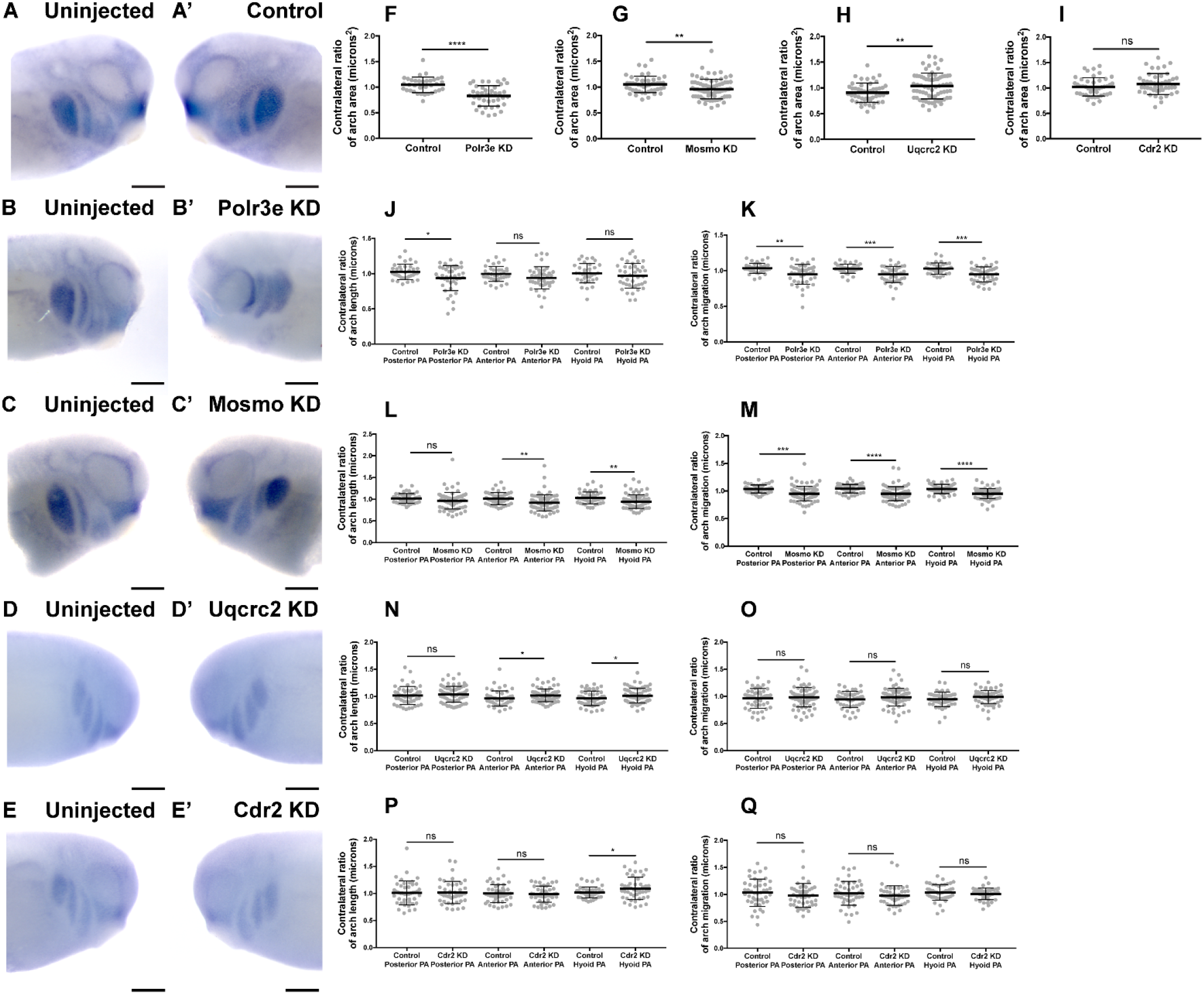
Knockdown of Polr3e and Mosmo affect CNC migration *in vivo*. **(A-C, A’-C) (A-E, A’-E’)** Lateral views of stage 25 embryos following whole-mount *in situ* hybridization against *twist*. Each column of panels are lateral views of two sides of the same embryo. Embryos were unilaterally injected to KD each individual 16p12.1-affected gene in half of the embryo and the other half was left uninjected. The left panels **(A-E)** represent the uninjected side and the right panels **(A’-E’)** represent the injected side. **(F-Q)** Measurements were taken for the total area of the three PAs (posterior PA, anterior PA, and hyoid PA), the length of each individual arch, and the NCC migration distance, as measured from the dorsal-most tip of each arch to the neural tube, by taking ratios of the injected side versus the uninjected side. Significance was determined using an unpaired students *t*-test with these ratios. **(F-H)** Partial depletion of both Polr3e and Mosmo significantly reduced the total area of the three PAs, while partial depletion of Uqcrc2 slightly increased the total area of the three PAs. **(J-K)** Polr3e KD significantly reduced the length of the posterior PA but had no effect on the length of the anterior PA or hyoid PA. However, partial depletion of this gene significantly reduced the total NCC migration distances for all three PAs. **(L-M)** Mosmo KD significantly reduced the length of the anterior PA and hyoid PA but had no effect on the length of the posterior PA. Partial depletion of this gene also significantly reduced the total NCC migration distance for all three PAs. **(N-O)** Uqcrc2 KD slightly increased the length of the anterior and hyoid PA but had no effect on the length of the posterior PA, nor did it affect the total NCC migration distance. **(I, P-Q)** Cdr2 KD had no effect on the total area of the three PAs, nor did it affect the total NCC migration distance. However, partial depletion of this gene slightly increased the length of the hyoid PA but did not affect the posterior or anterior PAs. (Embryos quantified: Control for Polr3e = 35, Control for Mosmo = 48, Control for Uqcrc2 = 48, Control for Cdr2 = 45, Polr3e KD = 41, Mosmo KD = 75, Uqcrc2 KD = 68, Cdr2 KD = 45). \*\*\*\**p* < 0.0001, \*\*\**p* < 0.001, \*\**p* < 0.01, \**p* < 0.05, n.s., not significant. Scalebar = 300μm.

While the previous experiment suggested a possible role for Polr3e, Mosmo, and Uqcrc2 in regulating NCC migration, other NCC-related defects might also lead to changes in measured PA area, length, and NCC migration distance. Thus, to investigate whether reduced dosage of any of the 16p12.1-affected genes impacted NCC migration rate itself, *in vitro* migration assays were performed, as previously described (Lasser et al., 2019; Mills et al., 2019). Individual genes were partially depleted in the whole embryo and NCCs were dissected prior to delamination from the neural tube (st. 17) from both KD and control conditions. These tissue explants were cultured on fibronectin-coated coverslips and individual cell migration for each explant was imaged using time-lapse phase-contrast microscopy for six hours (**Figures 4A-E,A’-E’**). Trajectories and speed of individual cells that escaped the explant were then mapped using automated particle tracking (**Figure 4F**) (Schindelin et al., 2012; Tinevez et al., 2017). Partial knockdown of Polr3e and Uqcrc2 resulted in slower individual NCC speed compared to controls (**Figures 4G,I**), while partial knockdown of Mosmo did not significantly impact NCC speed (**Figure 4H**). As partial knockdown of Cdr2 was not sufficient to significantly alter NCC streaming *in vivo*, nor was it sufficient to cause severe craniofacial or cartilage morphology defects, we hypothesized that NCC motility *in vitro* would not be affected by this depletion. Instead, knockdown of Cdr2 led to a significant increase in the speed of CNCs migrating *in vitro* (**Figure 4J**). We hypothesize that individual cell speed of Cdr2 KD *in vitro* may not directly correspond to differences in NCC streaming *in vivo* as the boundaries of NCC migration *in vivo* are heavily restricted due to repellent guidance cues within the PAs. Thus, together, we show that knockdown of Polr3e and Uqcrc2 alters NCC migration into the PAs, and that this defect could be driven by a reduction in individual NCC motility rates.

**Figure 4:**
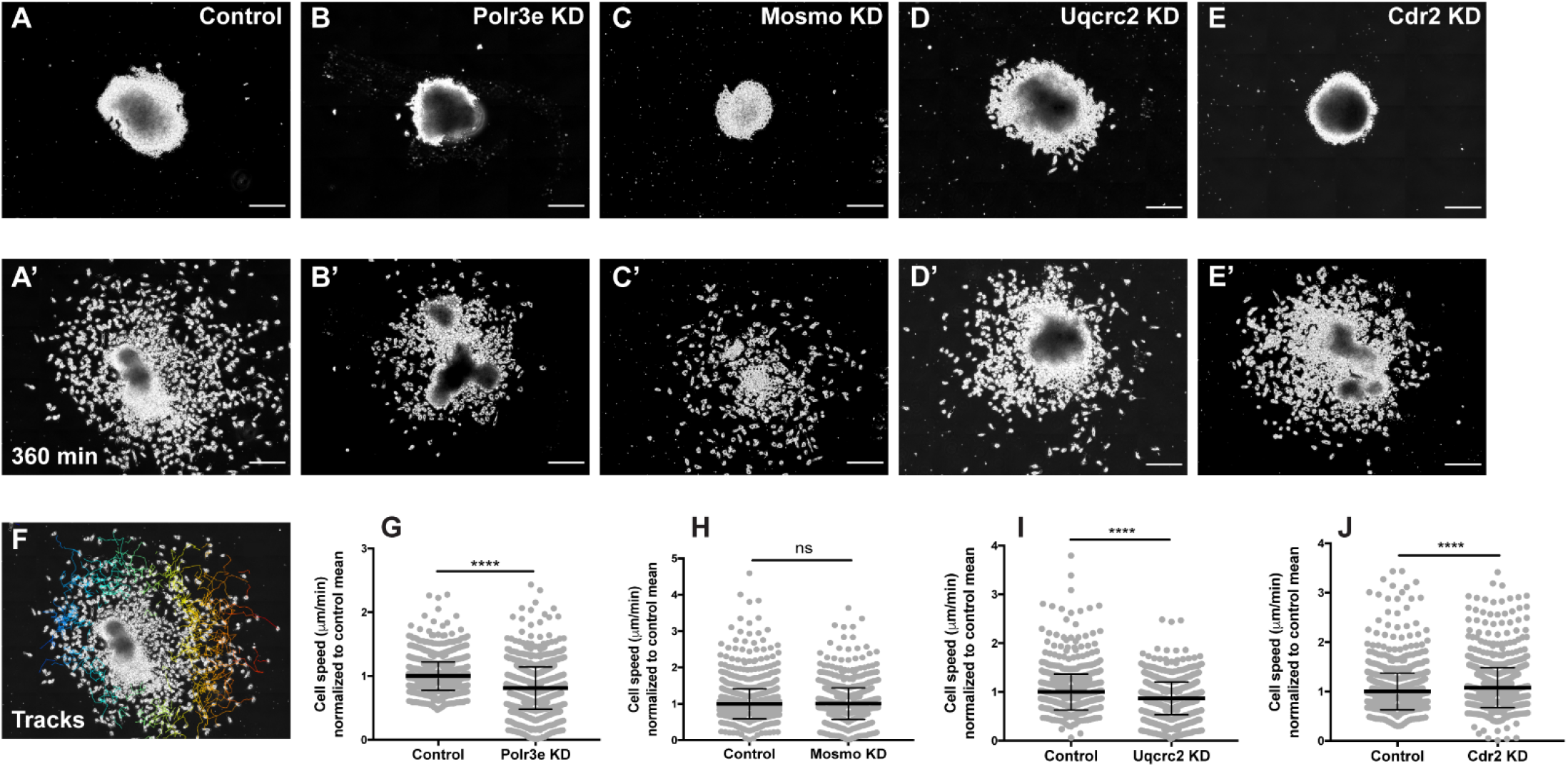
Manipulation of Polr3e, Uqcrc2, and Cdr2 impacts NCC migration speeds *in vitro*. Dissected NCC explants from control, Polr3e KD, Mosmo KD, Uqcrc2 KD, or Cdr2 KD embryos were plated on fibronectin-coated coverslips, allowed to adhere and begin migration, and imaged for 6 h using 20x phase microscopy. **(A-E)** Representative images of explants at initial timepoint (0 min). **(A’-E’)** Representative images of explants after 6 h migration (360 min). **(F)** Representative tracks generated by FiJi Trackmate plug-in. **(G-J)** Mean track speeds of Polr3e KD, Mosmo KD, Uqcrc2 KD, and Cdr2 KD explants compared to their controls. Partial depletion of Polr3e and Uqcrc2 significantly reduced mean NCC speed, while depletion of Cdr2 increased mean NCC speed. Partial depletion of Mosmo had no effect on mean NCC speed. (Explants quantified: 6-7 explants from control and KD embryos were plated for each experiment. Three separate experiments were performed for each depletion.) \*\*\*\**p* < 0.0001, \*\*\**p* < 0.001, \*\**p* < 0.01, \**p* < 0.05, n.s., not significant. Scalebar = 200μm.

### 16p12.1-affected genes do not directly impact NCC proliferation

As NCCs exit the dorsal neural tube and undergo directed migration along stereotypical pathways during vertebrate development, they must also balance between cell division and migration (Szabo and Mayor, 2018). Reduced gene dosage could result in smaller areas of *twist* expression due to decreased cellular migration rates, but it is also possible that there are fewer NCCs within the PAs if genetic manipulations affect NCC proliferation rates. To test this, each 16p12.1 gene was individually depleted in the whole embryo and NCCs were dissected as described in the previous section. These tissue explants were cultured on fibronectin-coated coverslips and NCCs were allowed to migrate away from the explant for four hours before being fixed. Immunocytochemistry was then performed using a phospho-histone H3 (PH3) antibody as a marker for cell proliferation (**Figures 5A-A’**). To measure NCC proliferation, the number of cells positively labeled for PH3 versus the total number of cells per explant were quantified using an automated particle counter after thresholding each image. Surprisingly, partial depletion of individual 16p12.1-affected genes had no statistically significant effect on NCC proliferation *in vitro* (**Figures 5B-E**). However, Mosmo KD resulted in a trend towards increased proliferation, and Cdr2 KD resulted in a trend towards decreased proliferation, suggesting a potential role for these genes in regulating this process (**Figures C,E**). Division of NCCs occurs over a wide range of times after exiting the neural tube, with mitotic activity significantly increasing as cells enter the pharyngeal arches (Gonsalvez et al., 2015; Rajan et al., 2018). Thus, it is possible that we do not observe a direct effect of genetic manipulation on NCC proliferation due to the absence of *in vivo* microenvironmental signals that are necessary for cell division.

**Figure 5:**
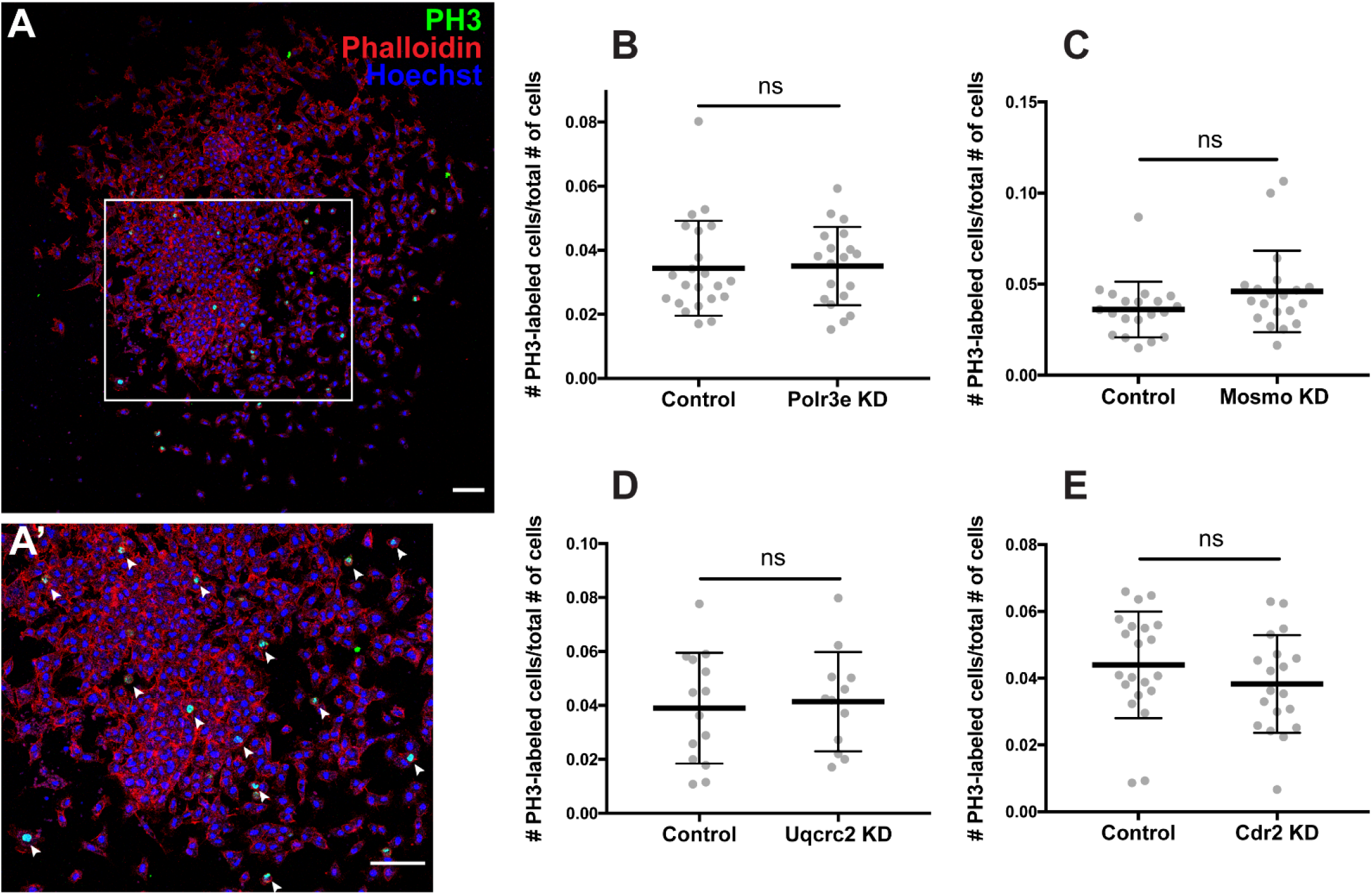
Manipulation of 16p12.1-affected genes does not impact NCC proliferation *in vitro*. Dissected NCC explants from control, Polr3e KD, Mosmo KD, Uqcrc2 KD, and Cdr2 KD embryos were plated on fibronectin-coated coverslips, allowed to adhere and begin migration for 4 h before being fixed in 4% PFA and immunostained with PH3 antibody as a marker for NCC proliferation, phalloidin to label the actin cytoskeleton, and Hoechst to label nuclei. **(A, A’)** Representative image of control NCC explant immunostained with PH3, phalloidin, and Hoechst. White arrows denote cells positively labeled for PH3. **(B-E)** Quantification of the number of positively PH3-labeled cells versus the total number of cells per NCC explant for Polr3e KD, Mosmo KD, Uqcrc2 KD, and Cdr2 KD compared to controls. Partial depletion of each individual 16p12.1-affected did not have a significant impact on NCC proliferation *in vitro*. (Explants quantified: 6-7 explants from control and KD embryos were plated for each experiment. Three separate experiments were performed for each depletion). \*\*\*\**p* < 0.0001, \*\*\**p* < 0.001, \*\**p* < 0.01, \**p* < 0.05, n.s., not significant. Scalebar = 300μm.

### Several 16p12.1-affected genes are critical for NCC induction and specification

The process of NCC induction and specification is complex and requires a specific level of signaling by the BMP, Wnt, FGF, RA, Shh, and Notch/Delta pathways to establish a gene regulatory network that is crucial for determining NCC identity (Pla and Monsoro-Burq, 2018; Prasad et al., 2019; Rogers and Nie, 2018; Theveneau and Mayor, 2012). During the early steps of NCC formation, these morphogen pathways work in concert with various NCC transcription factors (TFs) such as *snai1*, *snai2/slug*, *sox9*, and *twist*, to establish the neural plate border, regulate NCC specification, and subsequent NCC migration (Pla and Monsoro-Burq, 2018; Rogers and Nie, 2018). As shown in the previous sections, we found that partial depletion of several 16p12.1-affected genes significantly impacted *twist* expression patterns and NCC migration in the PAs, and that these defects were not due to changes in NCC proliferation rates. However, it is possible that reduced dosage of these genes causes NCC migration defects due to changes in NCC specification, resulting in fewer numbers of cells. Therefore, we tested this by utilizing single-blastomere injection strategies to generate left-right chimeric embryos, and compared side-by-side expression patterns of TFs required for NCC induction and specification between wild-type and KD sides.

Following single-sided individual depletion of each 16p12.1 gene, embryos were fixed at st. 16 and *in situ* hybridization was performed against *sox9* or *twist* (**Figures 6A-T)**. To quantify changes in expression of NCC specification markers, measurements were taken of the total area of expression of each marker for both the uninjected and control or KD side of each embryo (**Figures 6A-J**). Although both NCC specification markers were present, there were clear abnormalities including reduced signal and smaller total area of expression following partial knockdown of either Polr3e or Mosmo (**Figures 6C-F,K-L**). In contrast, knockdown of Uqcrc2 or Cdr2 did not significantly affect the signal or total area of expression for either NCC specification marker (**Figures 6G-L**). Together, these results suggest that Polr3e and Mosmo are distinctly required for NCC specification, as reduced levels alter expression of NCC specification markers, which may result in fewer cells and subsequent PA defects.

**Figure 6:**
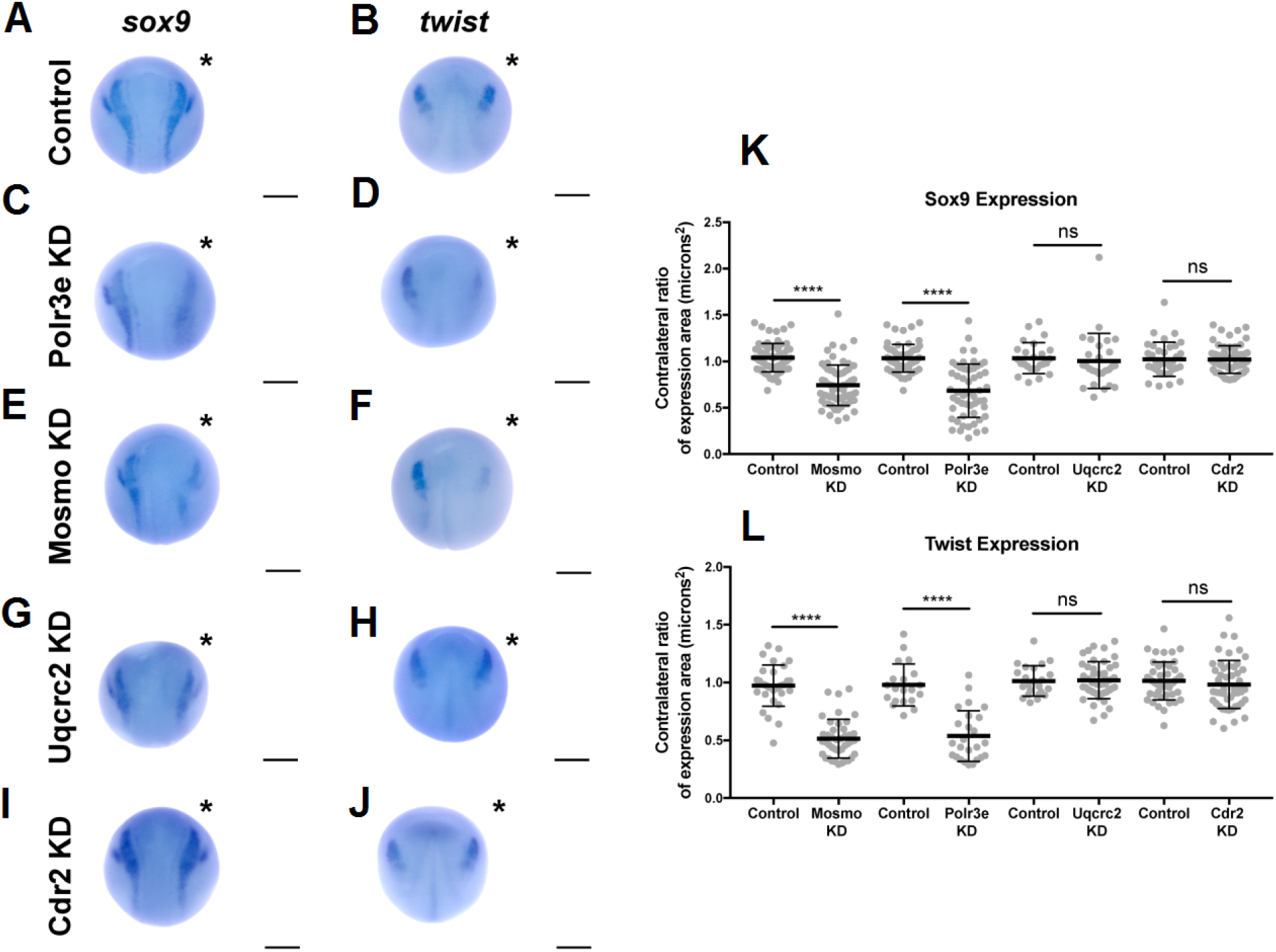
Manipulation of Polr3e and Mosmo affects NCC specification. *In situ* hybridization utilized **(A,C,E,G,I)** antisense mRNA probe against *sox9* and **(B,D,F,H,J)** antisense mRNA probe against *twist*. Each column of panels are dorsal views of two sides of the same embryo. Embryos were unilaterally injected to KD each individual 16p12.1-affected gene in half of the embryo and the other half was left uninjected. The left side represents the uninjected side and the right side, indicated with an asterisk (*), represents the injected side. Anterior of embryo is at the top. **(K-L)** Measurements were taken for the total area of the expression pattern for either *sox9* **(K)** or *twist* **(L)** using the polygon tool in ImageJ by taking ratios of the injected side versus the uninjected side. Significance was determined using an unpaired students *t*-test with these ratios. Partial depletion of either Mosmo or Polr3e significantly reduced the total area of expression for both *sox9* and *twist*, while partial depletion of either Uqcrc2 or Cdr2 did not significantly affect the total area of expression for either NCC specification marker. (Embryos quantified with *sox9* probe: Control for Polr3e = 60, Control for Mosmo = 66, Control for Uqcrc2 = 25, Control for Cdr2 = 36, Polr3e KD = 55, Mosmo KD = 69, Uqcrc2 KD = 27, Cdr2 KD = 59. Embryos quantified with *twist* probe: Control for Polr3e = 20, Control for Mosmo = 33, Control for Uqcrc2 = 23, Control for Cdr2 = 46, Polr3e KD = 25, Mosmo KD = 41, Uqcrc2 KD = 44, Cdr2 KD = 50). \*\*\*\**p* < 0.0001, \*\*\**p* < 0.001, \*\**p* < 0.01, \**p* < 0.05, n.s., not significant. Scalebar = 300μm.

## Discussion

To functionally explore the basis of the craniofacial and cartilage defects associated with the 16p12.1 deletion, we analyzed craniofacial phenotypes and cellular mechanisms underlying decreased dosage of four genes affected within this region. Our results show that three genes impacted by this CNV can contribute to normal craniofacial morphogenesis in *Xenopus laevis* (**Figure 7**), in a model where decreased gene dosage leads to both global and specific effects. We also provide evidence that deficits during NCC development may significantly contribute to the craniofacial dysmorphisms associated with the deletion. Specifically, we demonstrate, for the first time, that the 16p12.1-affected genes are co-expressed in motile NCCs and contribute to normal craniofacial patterning and cartilage formation. Several of these genes also directly impact NCC migration *in vivo* (*polr3e, mosmo*, and *uqcrc2*), NCC motility (*polr3e* and *uqcrc2*), and NCC specification (*polr3e* and *mosmo*), revealing new basic roles for these genes during embryonic development.

**Figure 7:**
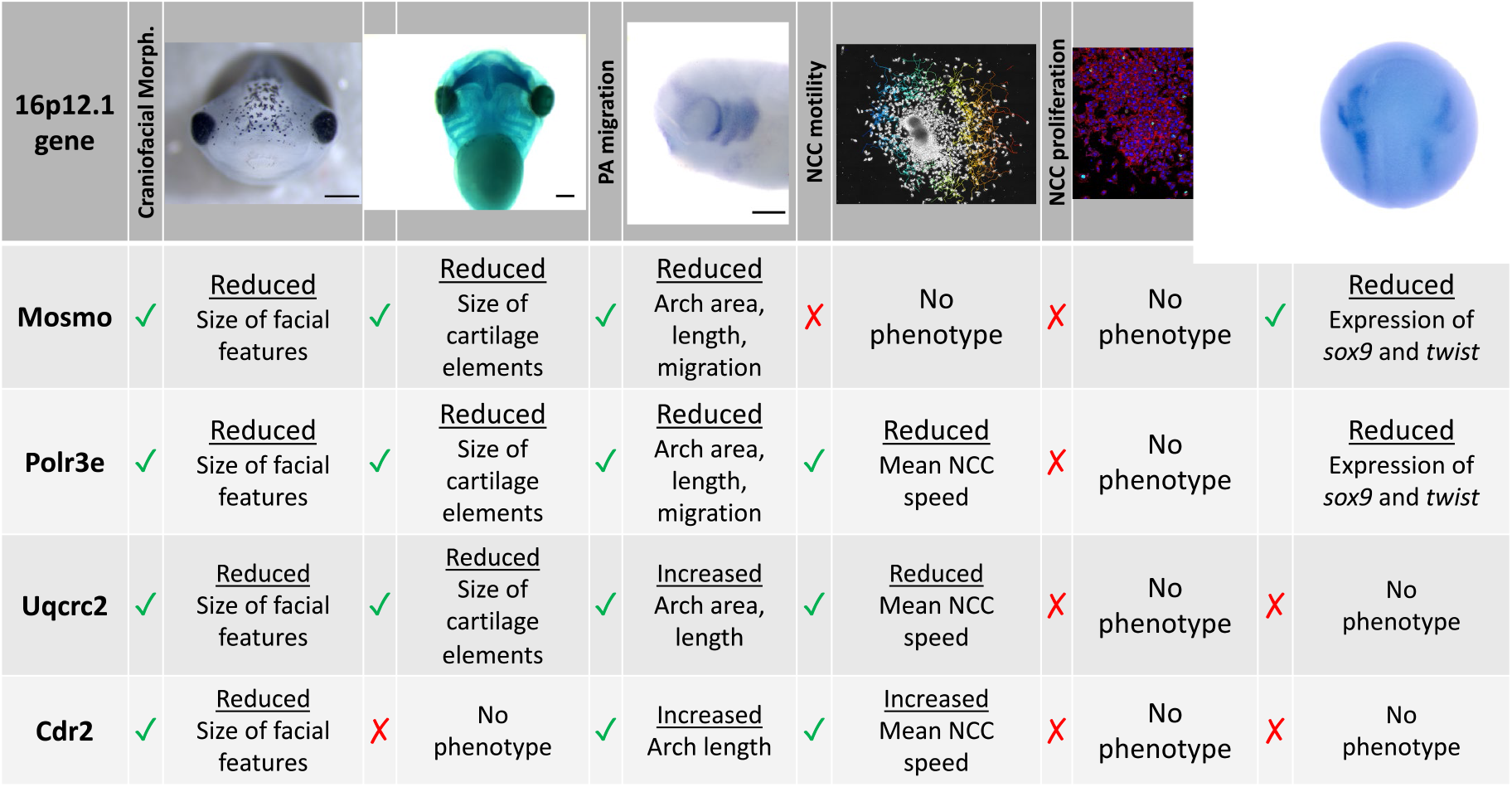
Summary table of 16p12.1-affected gene craniofacial, cartilage, and NCC phenotypes. Partial depletion of 16p12.1-affected genes demonstrates numerous impacts on craniofacial, cartilage, and CNC development. Tissues are denoted as affected (checked box) if phenotypes were significantly different from control (p < 0.05); see individual figures for data distribution and statistics.

As the 16p12.1 deletion is multigenic in nature, we sought to determine each gene’s contribution to a craniofacial phenotype. Although we have narrowed our studies to focus on how each individual 16p12.1-affected gene contributes to facial patterning, our findings align well with the idea that the presentation of symptoms associated with the deletion is a cumulative product of the impacted region. While *polr3e* and *mosmo* knockdowns severely impacted nearly all examined aspects of craniofacial morphogenesis and NCC development across early developmental stages, *uqcrc2* and *cdr2* knockdowns produced very minimal or no phenotypes in these areas.

In particular, we observed a global functional role for *polr3e*, *mosmo*, and *uqcrc2* in both craniofacial and cartilage development. Individual depletion of these genes narrowed facial width and area, and decreased eye size in a way that appears analogous to the smaller head and smaller eye size phenotypes observed in children with the 16p12.1 deletion (Pizzo et al., 2020). Moreover, reduced levels of these genes decreased the size of cartilaginous tissue structures important for jaw and mouth formation, which may correlate with the micrognathia phenotype. However, the phenotypes associated with *uqcrc2* KD were quite mild in comparison to *polr3e* or *mosmo* KD, and *cdr2* KD had little to no effect on craniofacial or cartilage formation. Thus, further investigation into how these gene depletions function combinatorially to generate the full signature of the 16p12.1 deletion craniofacial dysmorphism is necessary.

It is worthwhile to mention that depletion of the 16p12.1-affected genes in *X. laevis* almost certainly diverges from perfect recapitulation of the 16p12.1 deletion symptoms. *Xenopus* has emerged as a powerful model system to study human genetic diseases of craniofacial development, as the majority of disease-associated pathways that drive craniofacial morphogenesis are conserved between these species (Devotta et al., 2018; Dickinson, 2016; Dubey and Saint-Jeannet, 2017; Griffin et al., 2018; Lasser et al., 2019; Lichtig et al., 2020; Mills et al., 2019; Schwenty-Lara et al., 2020; Tahir et al., 2014). However, not only does the CNV contain non-coding regions that we did not model here, but there are also some craniofacial and cartilage morphological differences between *Xenopus* and humans that prevent certain direct correlations to disease pathology. For example, NCCs residing in the hyoid PA, that give rise to the ceratohyal cartilage in *Xenopus,* will later give rise to anterior portions of the face and combine with contributions from the Meckel’s cartilage to form regions of the lower jaw (Gross and Hanken, 2008; Kerney et al., 2012). In humans, Meckel’s cartilage will similarly form portions of the lower jaw but will also become part of the middle ear skeletal structures. Therefore, morphological impacts resulting from aberrant development of these tissues may have more direct correlates to human pathology in the context of NCC migration and development of the PAs.

Within that effort, our work has demonstrated co-expression of these gene transcripts in NCCs and their necessity during specific NCC-related processe,s which may be a driving mechanism underlying the observed craniofacial and cartilage defects. Decreased dosage of *polr3e* or *mosmo* significantly affected PA migration *in vivo* leading to decreased PA area, length, and NCC migration distance from the neural tube. Although *uqcrc2* KD led to both craniofacial and cartilage defects, surprisingly, its depletion caused an increase in PA area and length, possibly due to inappropriate proliferation and expansion of NCCs. Our data suggests that these PA migration defects are likely due to gene-specific effects during aspects of NCC development, as *polr3e* KD impacted both NCC motility and specification, *mosmo* KD impacted NCC specification, and *uqcrc2* KD impacted NCC motility.

While we show that several 16p12.1-affected genes are important for regulating NCC development and subsequent formation of their tissue derivatives, it remains unclear as to why any of these genes may be exceptionally critical in these tissues. This question must be left to some speculation, as the cell biological functions of these genes are extremely diverse and warrant further comprehensive investigation in the context of embryonic craniofacial and NCC development. However, a brief summary of the known roles of these genes and how they may mechanistically influence NCC-related processes are outlined here.

POLR3E is a subunit of RNA polymerase III, important for regulating the transcription of small RNAs, such as 5S rRNA and tRNAs (Hu et al., 2002). However, the precise role of this subunit in relation to RNA polymerase III activity has not been well studied. It is known that the function of RNA polymerase III is tightly dependent on cell growth and differentiation, supporting the idea that its alteration would lead to deficits in these processes (Dumay-Odelot et al., 2010). While depletion of *polr3e* did not significantly alter NCC proliferation, it did affect NCC specification, suggesting that it may be important for regulating transcription of genes necessary in maintaining the identity and subsequent differentiation of NCCs. Moreover, mutations in other subunits of RNA polymerase III, POLR1C and POLR1D, have been associated with leukodystrophy, ataxia, and the congenital craniofacial disorder, Treacher Collins syndrome (Ghesh et al., 2019; Kadakia et al., 2014; Noack Watt et al., 2016; Papageorgiou et al., 2020). Studies suggest that loss-of-function of these genes results in cartilage hypoplasia and cranioskeletal anomalies, due to deficient ribosome biogenesis, increases in cellular death, and deficiencies in NCC migration (Noack Watt et al., 2016). Given that POLR3E has been shown to interact with POLR1C and POLR1D, it is plausible that it may function in a similar capacity.

Recent work in cell culture suggests that MOSMO acts as a negative regulator of Sonic hedgehog signaling (Shh) by degrading the Frizzled class receptor, Smoothened (Pusapati et al., 2018), and our work is the first to elucidate the *in vivo* cellular function of this gene in the context of vertebrate embryonic craniofacial development. The Shh pathway is known to be critical for craniofacial morphogenesis, as upregulation or downregulation of signaling can lead to aberrant NCC patterning, development, and maintenance (Abramyan, 2019; da Costa et al., 2018; Dworkin et al., 2016; Everson et al., 2017; Grieco and Hlusko, 2016; Hammond et al., 2018; Millington et al., 2017; Okuhara et al., 2019; Wang et al., 2019). Shh signaling coordinates the downstream intracellular activity of Gli TFs, which stimulate transcription of several target genes required for NCC induction and specification (Cerrizuela et al., 2018; Millington et al., 2017; Rogers and Nie, 2018). This aligns well with our results demonstrating that *mosmo* can directly impact NCC specification, highlighting a new cell biological role for this gene. Additionally, Shh interacts with other morphogenic signaling pathways, like BMP, Wnt, FGF, and Notch, all of which are required for NCC development. In particular, enhancement of the Shh gradient can restrict canonical Wnt signaling by promoting expression of genes encoding Wnt inhibitors, causing an increase in NCC proliferation and expansion that eventually results in craniofacial defects such as cleft lip (Hammond et al., 2018; Kurosaka et al., 2014). As MOSMO is a negative regulator of Shh signaling, its depletion should lead to upregulation of the pathway. Therefore, it is possible that reducing *mosmo* dosage disrupts downstream target gene expression, specifically genes required for NCC specification, and perturbs Wnt signaling, such that NCC development is impacted, leading to craniofacial and cartilage morphogenesis defects.

UQCRC2 is a component of the mitochondrial respiratory chain complex III that is required for its assembly and is important for normal mitochondrial activity to produce ATP (Gaignard et al., 2017; Hammond et al., 2018; Kurosaka et al., 2014; Miyake et al., 2013). Reduced levels of *uqcrc2* produced both craniofacial and cartilage phenotypes, potentially due to decreased NCC motility. This could be especially damaging in the context of multipotent NCCs, as metabolism is increasingly demonstrated to perform a commanding role in determination of cell fate and subsequent motility (Mathieu and Ruohola-Baker, 2017; Perestrelo et al., 2018; Sperber et al., 2015). Our results also demonstrate that depletion of *uqcrc2* leads to an increase in PA area and length, potentially due to unchecked NCC expansion and proliferation. Studies suggest that overexpression of UQCRC2 is correlated with tumor progression through increased cellular proliferation (Shang et al., 2018). Thus, its depletion may also upregulate cell proliferation through a feedback mechanism to compensate for lower gene dosage levels. Moreover, patients with mutations of this gene have recurrent liver failure, lactic acidosis, and hypoglycemia (Gaignard et al., 2017; Miyake et al., 2013). As *Xenopus* is an excellent model for studying the development of both liver and kidney organ systems (Blackburn et al., 2019; Blackburn and Miller, 2019), and as we have shown expression of *uqcrc2* in the *Xenopus* kidney, it would be interesting to further examine how this gene mechanistically regulates development and function of these other tissues.

CDR2 is an oncogenic protein that is strongly expressed in the cerebellum and is ectopically produced by tumor cells, specifically in ovarian and breast malignancies (Hwang et al., 2016; Schubert et al., 2014; Venkatraman and Opal, 2016). Loss of immune tolerance to this protein is believed to trigger the synthesis of the autoantibody, leading to an immunological reaction causing cerebellar degeneration. CDR2 is also believed to play multiple roles in the regulation of transcription by interacting with the TF c-Myc, sequestering it in the cytoplasm, and inhibiting its transcription of downstream target genes (O’Donovan et al., 2010; Okano et al., 1999). It is known that the expression of c-Myc is required for correct temporal and spatial development of NCCs (Bellmeyer et al., 2003), and while, here, we focused on exploring how depletion of *cdr2* affects craniofacial morphogenesis, understanding how overexpression of this gene affects NCC development warrants further study. *Cdr2* mRNA can be found in almost all cell types, however, its protein expression is limited to Purkinje neurons, some brainstem areas, and spermatogonia (Venkatraman and Opal, 2016). This could explain why depletion of this gene in *Xenopus* did not produce significant craniofacial or cartilage defects, nor affect NCC developmental processes. It is also possible that other closely related genes, such as *cdr1*, *cdr3*, or *cdr2l*, can functionally compensate for the lack of *cdr2* levels. Altogether, it is clear that our current knowledge of how these genes ultimately contribute to embryonic craniofacial and cartilage morphogenesis is lacking, and further basic cell biological examination of 16p12.1-affected gene function within a developmental context is necessary for a better mechanistic understanding of the 16p12.1 deletion etiology.

Finally, it will also be essential to explore how these genes ultimately synergistically or epistatically regulate the pathology associated with the 16p12.1 deletion. To this aim, our model provides a unique advantage by being a moderate throughput, titratable, and inexpensive system to combinatorially deplete numerous genes simultaneously. Together, our current and ongoing work suggests significant roles for several 16p12.1-affected genes as potent effectors of NCC-derived tissues that regulate specific processes during their development, providing a foundation for the underlying mechanisms contributing to the craniofacial defects associated with the 16p12.1 deletion.

## Materials and Methods

### *Xenopus* husbandry

Eggs obtained from female *Xenopus laevis* were fertilized in vitro, dejellied and cultured at 14-23°C in 0.X Marc’s modified Ringer’s (MMR) using standard methods (Sive et al., 2010). Embryos received injections of exogenous mRNAs or antisense oligonucleotide strategies at the two or four cell stage, using four total injections performed in 0.1X MMR media containing 5% Ficoll. Embryos were staged according to Nieuwkoop and Faber (Nieuwkoop and Faber, 1994). All experiments were approved by the Boston College and Boston University Institutional Animal Care and Use Committees and were performed according to national regulatory standards.

### Depletion and Rescue

Morpholinos (MOs) were targeted to early splice sites of *X. laevis mosmo* (for L, 5-ACAATTGACATCCACTTACTGCCGG-3; for S, 5-CACCTTCCCTACCCCGCTACTTAC-3), *polr3e* (for L, 5-ACTGTAAGCCTCTTTTGCCTTACCT-3), *uqcrc2* (for L, 5-ACAGTGTCTCTAAAGCACAGATACA-3; for S, 5-CCCCTAACCCATTAAACATATACCT-3), *cdr2* (for L and S, 5-CATCCCTCCCATACACTCACCTTG-3), or standard control MO (5-cctcttacctcagttacaatttata-3); purchased from Gene Tools (Philomath, OR). In knockdown (KD) experiments, all MOs were injected at either the 2-cell or 4-cell stage with embryos receiving injections 2 or 4 times total. *Mosmo* and control MOs were injected at 12ng/embryo for 50% KD and 20ng/embryo for 80% KD; *polr3e* and control MOs were injected at 5ng/embryo for 30% KD, 10ng/embryo for 50% KD and 20ng/embryo for 80% KD; *uqcrc2* and control MOs were injected at 35ng/embryo for 50% KD and 50ng/embryo for 80% KD; *cdr2* and control MOs were injected at 10ng/embryo for 50% KD. Splice site MOs were validated through a Reverse Transcriptase Polymerase Chain Reaction (RT-PCR) as previously described (Pizzo et al., 2020).

Rescues were performed with exogenous mRNAs co-injected with their corresponding MO strategies (Pizzo et al., 2020). *Xenopus* ORFs for *mosmo*, *polr3e*, and *uqcrc2* were purchased from the European *Xenopus* Resource Center (EXRC) and gateway-cloned into pCSF107mT-GATEWAY-3’GFP destination vectors. Constructs used were *mosmo*-*gfp*, *polr3e*-*gfp*, *uqcrc2*-*gfp*, *cdr2*-*gfp*, and *gfp* in pCS2+. In rescue experiments, MOs (amounts used as in KD for each MO) were injected with mRNA (1500pg/embryo for *mosmo-gfp*; 2000pg/embryo for *polr3e*-*gfp*; 1000pg/embryo for *uqcrc2-gfp*; 1000pg/embryo for *cdr2-gfp*) in the same injection solution.

### Whole-mount RNA *in situ* hybridization

Embryos were fixed overnight at 4°C in a solution of 4% paraformaldehyde in PBS, gradually dehydrated in ascending concentration of methanol in PBS, and stored in methanol at −20°C for a minimum of two hours, before *in situ* hybridization, performed as previously described (Sive et al., 2007). After brief proteinase K treatment, embryos were bleached under a fluorescent light in 1.8x saline-sodium citrate, 1.5% H2O2, and 5% (vol/vol) formamide for 20 minutes to 45 minutes before prehybridization. During hybridization, probe concentration was 0.5ug/mL.

The *Xenopus sox9* hybridization probe was a kind gift from Dr. Dominique Alfandari (University of Massachusetts at Amherst, MA), and the *Xenopus twist* hybridization probe was a kind gift from Dr. Richard Harland and Dr. Helen Willsey (University of California Berkeley and University of California SF, CA). The templates for making antisense probes for *polr3e*, *mosmo*, *uqcrc2*, and *cdr2* were PCR amplified from the reverse transcribed cDNA library, using the following primer sets: *polr3e* forward, 5’ – GGATAGTCGCTCAGAACACG – 3’, *polr3e* reverse, 5’ – GGGTCAGCTTTGTCTGGATC – 3’, *mosmo* forward, 5’ – TCTGGATGTTTGTTTCTGGCTGC – 3’, *mosmo* reverse, 5’ – GGGTAATTTGTAGGGTTGGCCTC – 3’, *uqcrc2* forward, 5’ – TCCTCTCTAGGAGGCTTTACTCTG – 3’, *uqcrc2* reverse, 5’ – GGGAGCCAATTTCACCAATCAG – 3’, *cdr2* forward, 5’ – GACAGCAACGTGGAGGAGTTC – 3’, and *cdr2* reverse, 5’ – GCGCAGATCATACAGCTCCTTC – 3’. The antisense digoxigenin-labeled hybridization probes were transcribed *in vitro* using the T7 MAXIscript kit. Embryos were imaged using a Zeiss AxioCam MRc attached to a Zeiss SteREO Discovery.V8 light microscope. Images were processed in ImageJ.

### Cartilage staining

At stage 42, *Xenopus* embryos were anesthetized with benzocaine and fixed in 4% paraformaldehyde in PBS overnight. Alcian blue staining of embryos was performed based on the Harland Lab protocol. Before ethanol dehydration, embryos were bleached under a fluorescent light in 1.8x saline-sodium citrate, 1.5 H2O2, and 5% (vol/vol) formamide for 30 minutes. Embryos were imaged in PBS, using a Zeiss AxioCam MRc attached to a Zeiss SteREO Discovery.V8 light microscope. Images were processed in ImageJ. Analysis of cartilage structures was performed in ImageJ utilizing the polygon, area, and line functions. Measurements included the average ceratohyal area (outlined cartilage in Fig 2), and the branchial arch width, which was quantified by taking the width of the branchial arch across the widest point. Differences were analyzed by student’s unpaired t-test using Graphpad (Prism).

### Half-embryo injections

Half KDs were performed at the two-cell stage. *X. laevis* embryos were unilaterally injected two times with either control MO or 16p12.1 gene-specific MO and a *gfp* mRNA construct. The other blastomere was left uninjected. Embryos were raised in 0.1X MMR through neurulation, then sorted based on left/right fluorescence. For NCC specification experiments, embryos were fixed at stage 16, and for pharyngeal arch (PA) visualization, embryos were fixed between stage 25-30. Whole-mount *in situ* hybridization was then performed according to the previously described procedure. Analysis of NCC specification markers from *in situ* experiments was performed on dorsal view images in ImageJ by measuring the total area of expression using the polygon tool. Analysis of PAs from *in situ* experiments was performed on lateral view images in ImageJ. Measurements were taken to acquire: 1) arch area, the area of individual PA determined using the polygon tool, 2) arch length, the length of the distance between the top and bottom of each PA, and 3) arch migration, the ventral most part of the PA to the neural tube. All measurements were quantified by taking the ratio between the injected side versus the uninjected side for each sample, respectively. Statistical significance was determined using a student’s unpaired t-test in Graphpad (Prism).

### Neural crest explants, imaging, and analysis

Embryos at stage 17 were placed in modified DFA solution (53mM NaCl, 11.7mM NA2CO3, 4.25mM K Gluc, 2mM MgSO4, 1mM CaCl2, 17.5 mM Bicine, with 50ug/mL Gentamycin Sulfate, pH 8.3), before being stripped of vitelline membranes and imbedded in clay with the anterior dorsal regions exposed. Skin was removed above the NCC using an eyelash knife, and NCCs were excised. Explants were rinsed, and plated on fibronectin-coated coverslips in imaging chambers filled with fresh DFA. Tissues were allowed to adhere forty-five minutes before being moved to the microscope for time-lapse imaging of NCC motility. Microscopy was performed on a Zeiss Axio Observer inverted motorized microscope with a Zeiss 20x N-Achroplan 0.45 NA Phase-contrast lens, using a Zeiss AxioCam camera controlled with Zen software. Images were collected using large tiled acquisitions to capture the entire migratory field. Eight to ten explants, from both control and experimental conditions, were imaged at a six-minute interval, for six hours. Data was imported to ImageJ, background subtracted, and cropped to a uniform field size. Migration tracks of individual cells were collected manually using the Manual Tracking plug-in. Mean speed rates were imported to Graphpad (Prism), and compared between conditions using student’s unpaired t-tests. Three independent experiments were performed for each condition.

For NCC proliferation, NCC tissue explants were allowed to adhere and migrate on fibronectin-coated coverslips for four hours before being fixed in 4% paraformaldehyde. Explants were permeabilized in 0.1% Triton-X 100 in PBS, blocked with a solution containing 2% bovine serum albumin, 0.1% Triton-X 100 in PBS, and incubated in Phospho-Histone H3 (Ser10) (Invitrogen, PA5-17869, polyclonal, 1:500), goat anti-rabbit Alexa Fluor^488^ conjugate secondary antibody (Invitrogen, 1:1000), Alexa Fluor^568^ phalloidin (Invitrogen, 1:500), and Hoechst 33342 solution (Invitrogen, 1:1000). Microscopy was performed on a Zeiss AiryScan inverted motorized microscope with a Zeiss 20X lens, using a Zeiss AxioCam camera controlled with Zen software. Images were acquired using large tiled acquisitions to capture the entire migratory field. Images of five to seven explants, from both control and experimental conditions were imported to ImageJ, and the total number of PH3-labeled positive cells versus the total number of cells were quantified using an automated particle counter after thresholding each image. Cell counts were imported to Graphpad (Prism), and compared between conditions using student’s unpaired t-test. Three independent experiments were performed for each condition.

## Acknowledgements

We thank Helen Willsey for reading and comments on the manuscript. We thank members of the Lowery Lab for helpful discussions, suggestions, and editing. We thank Connor Monahan and Sydney Kim for technical assistance. We thank Nancy McGilloway and Todd Gaines for excellent *Xenopus* husbandry. We also thank the National *Xenopus* Resource (RRID:SCR013731) and Xenbase (RRID:SCR-003280) for their support. We thank Bret Judson and the Boston College Imaging Core for infrastructure and support. This work was supported by the National Institutes of Health [R01 GM121907, R01 MH109651, R03 DE025824], the American Cancer Society [RSG-16–144-01-CSM], and National Science Foundation [Grant No. 1626072].

---

**Figure S1:**
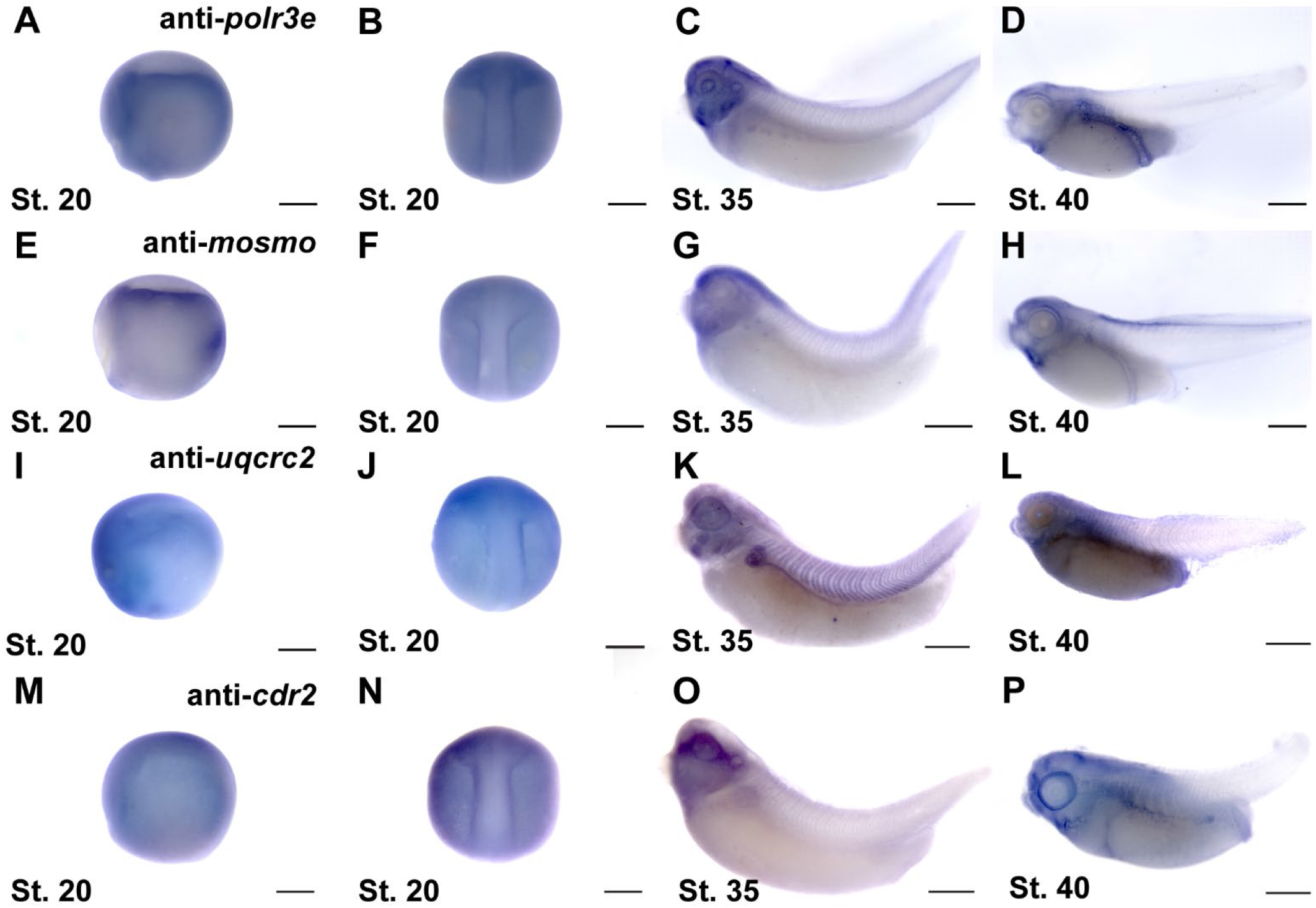
Expression patterns for 16p12.1-affected genes across early development. *In situ* hybridization utilized **(A-D)** antisense mRNA probe to *polr3e*, **(E-H)** antisense mRNA probe to *mosmo*, **(I-L)** antisense mRNA probe to *uqcrc2*, and **(M-P)** antisense mRNA probe to *cdr2*. Anterior to the left. Lateral and dorsal view images of embryos shown at stage 20 **(A-B, E-F, I-J, M-N)**, lateral view at stage 35 **(C, G, K, O)**, and lateral view at stage 40 **(D, H, L, P)** (n = 10 per probe) Scalebar = 300μm.

**Figure S2:**
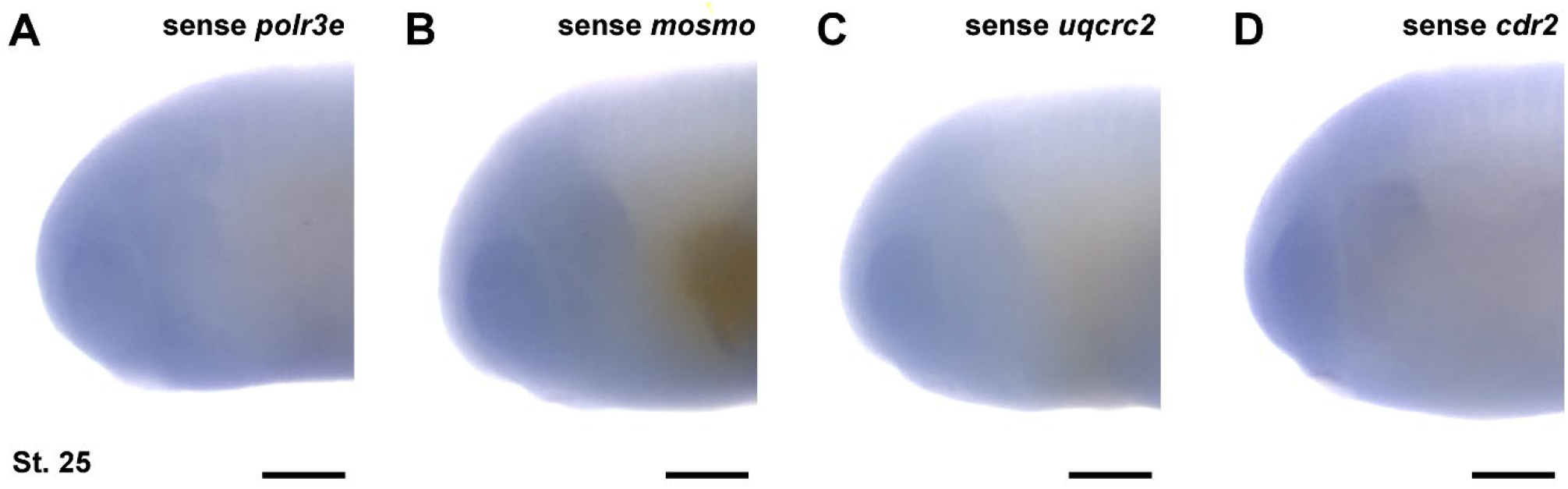
*In situ* hybridization probes generated against sense strands of 16p12.1-affected gene mRNAs. In situ hybridization utilized **(A)** sense mRNA probe against *polr3e*, **(B)** sense mRNA probe against *mosmo*, **(C)** sense mRNA probe against *uqcrc2*, and **(D)** sense mRNA probe against *cdr2*, shown at stage 25. (n = 10 per probe). Scalebar = 300μm.

